# Molecular Architecture of the Bardet-Biedl Syndrome Protein 2-7-9 Subcomplex

**DOI:** 10.1101/699223

**Authors:** W. Grant Ludlam, Takuma Aoba, Jorge Cuéllar, M. Teresa Bueno-Carrasco, Aman Makaju, James D. Moody, Sarah Franklin, José M. Valpuesta, Barry M. Willardson

**Affiliations:** Department of Chemistry and Biochemistry, Brigham Young University, Provo, UT 84602; Centro Nacional de Biotecnología (CNB-CSIC), Campus de la Universidad Autónoma de Madrid, 28049 Madrid, Spain; Department of Internal Medicine, Nora Eccles Harrison Cardiovascular Research and Training Institute, University of Utah, Salt Lake City, UT 84112

**Keywords:** Bardet-Biedl Syndrome, Ciliopathy, Integrated modeling, cilia transport, protein assembly, protein complex, protein cross-linking, mass spectrometry (MS), electron microsopy (EM), homology modeling

## Abstract

Bardet-Biedl syndrome (BBS) is a genetic disease caused by mutations that disrupt the function of the BBSome, an eight-subunit complex that plays an important role in transport of proteins in primary cilia. To better understand the molecular basis of the disease, we analyzed the structure of a BBSome subcomplex consisting of three homologous BBS proteins (BBS2, BBS7, and BBS9) by an integrative structural modeling approach using electron microscopy and chemical crosslinking coupled with mass spectrometry. The resulting molecular model revealed an overall structure that resembles a flattened triangle. Within the structure, BBS2 and BBS7 form a tight dimer based on a coiled-coil interaction, and BBS9 associates with the dimer via an interaction with the α-helical domain of BBS2. Interestingly, a BBS-linked mutation of BBS2 (R632P) is located in the α-helical domain at the interface between BBS2 and BBS9, and binding experiments showed that this mutation disrupted the interaction of BBS2 with BBS9. This finding suggests that BBSome assembly is disrupted by the R632P substitution, providing a molecular explanation for BBS in patients harboring this mutation.

## Introduction

Primary cilia perform vital signaling functions in vertebrate cells, ranging from recognition of developmental cues from morphogens such as hedgehog in the developing embryo to detection of sensory signals such as photons of light in retinal photoreceptor cells (1–4). Primary cilia are formed by the axoneme, a circularly-ordered scaffold containing nine pairs of microtubules anchored inside the cell at the basal body and protruding outward to create a finger-like projection of the plasma membrane (1, 2). Many transmembrane receptors are concentrated in this ciliary compartment, creating a type of signaling antenna for the cell (5–7). The contents of the cilium are delivered there by the intraflagellar transport (IFT) complexes that use kinesin motors to move cargos toward the tip of the cilium (anterograde transport) and dynein motors to move cargos toward the base of the cilium (retrograde transport) (1,2,4). Failure of ciliary trafficking results in diseases referred to as ciliopathies that are characterized by multiple phenotypes, including cystic kidneys, retinal degeneration, obesity and multiple developmental disorders (8).

One of these ciliopathies is Bardet-Biedl syndrome (BBS), a disease that results from the malfunction of a large protein complex called the BBSome (9). The BBSome consists of eight subunits (named BBS1, 2, 4, 5, 7, 8, 9 and 18), and mutations in each subunit have been linked to BBS (9–12). The disease results from an inability of the BBSome to participate in ciliary transport. The proposed function of the BBSome is to act as a scaffolding complex to link protein cargos, including membrane proteins, to the IFT machinery for ciliary transport with a particularly important role in retrograde transport out of the cilium (11,13–18). The BBSome reversibly associates with the membrane via an interaction of BBS1 with ARL6 (ADP-ribosylation factor-like protein 6), a small GTPase that interacts with the membrane in its GTP-bound form (11, 19). In this membrane-bound state, the BBSome is believed to pick up membrane proteins targeted for exit from the cilium (11, 18). Transport of these cargos occurs through association of the BBSome with the IFT-B complex via an interaction with a linker protein LZTFL1 (Leucine zipper transcription factor-like 1) that binds to IFT27 in the IFT-B complex (15,16,20). In this manner, the BBSome collects membrane proteins for IFT- and dynein-mediated retrograde transport.

To perform its transport function, each of the eight subunits of the BBSome must be translated on ribosomes, folded into their native state and assembled into a functional complex. Evidence suggests that BBSome assembly proceeds through several subcomplexes, with BBS9 acting as a central scaffold. A stable hexameric complex consisting of human BBS1, 4, 5, 8, 9 and 18 was recently isolated from insect cells (21). This subcomplex interacted with Arl6 as well as peptide motifs from several G protein-coupled receptor cargos. The two other subunits of the BBSome, BBS2 and BBS7, have an intricate folding and assembly process that requires a network of molecular chaperones, including the cytosolic chaperonin containing tailless polypeptide 1 (CCT; also termed TRiC) complex and three chaperonin-like proteins named BBS6, 10 and 12 (22, 23). Inactivating mutations in these chaperonin-like BBS proteins are also a major cause of BBS (24–26). The chaperonin proteins associate with BBS2 and BBS7 and assist in the formation of a BBS2/BBS7 dimer that binds BBS9 via an interaction with BBS2 (23, 27). Presumably, binding of the BBS2/BBS7 dimer with the hexameric complex completes the assembly of the BBSome octamer (21).

To better understand the mechanism of BBSome assembly, its function as a scaffold for intraflagellar transport and the molecular basis of BBS disease, we have isolated a trimeric BBSome subcomplex of BBS2, BBS7 and BBS9 and investigated its structure by electron microscopy (EM) and chemical crosslinking coupled with mass spectrometry (XL-MS) using the integrative modeling platform (28). The data show that the BBS2/BBS7 dimer is stabilized by an extensive coiled-coil interaction and that BBS9 interacts with the dimer through association with the α-helical domain of BBS2. A BBS-causing mutation in BBS2 (R632P) (29–31) in this region disrupts the interaction of BBS2 with BBS9, suggesting that the inability of the BBS2/BBS7 dimer to associate with the hexameric complex is the underlying cause of BBS in patients with the R632P mutation.

## Results

### BBS2-7-9 purification

The complex between BBS2, BBS7 and BBS9 is an important early intermediate in assembly of the BBSome (23). We purified this BBS2-7-9 complex for structural studies by co-expressing affinity-tagged versions of each subunit in HEK-293T cells and isolating complexes containing the Strep peptide-tagged BBS7 using a Strep-Tactin column. This purification resulted in roughly equal amounts of BBS2, BBS7, BBS9 with a 70 kDa contaminant protein corresponding to heat shock protein 70 (Hsp70) isoforms (Fig. 1A). The complex was further purified by glycerol gradient centrifugation (Fig. 1B), which separated Hsp70 and other contaminants (fractions 3-9 and 15-25) from the core complex (fractions 10-13).

**Figure 1.**
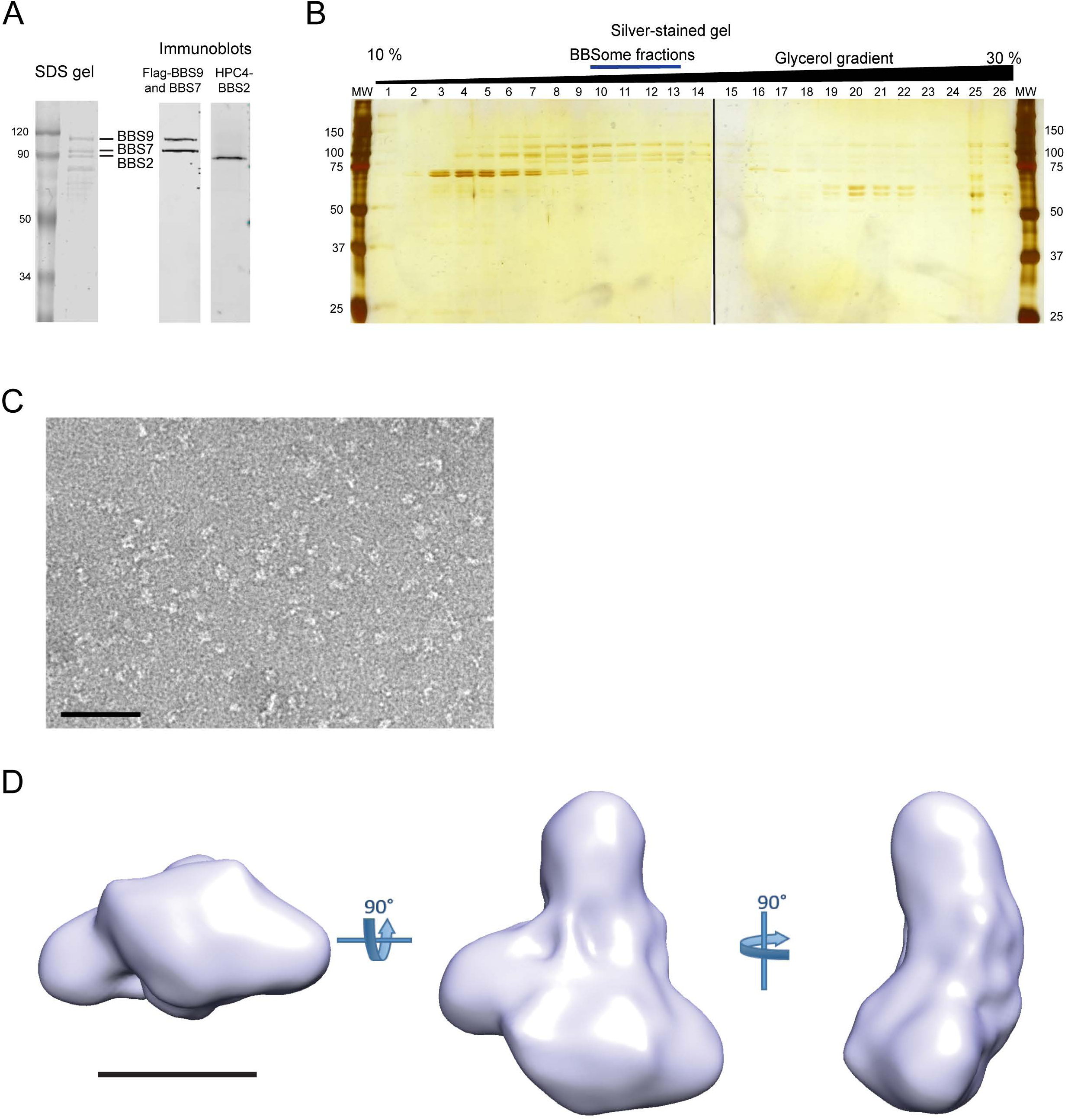
Purification of the BBS2-7-9 Complex and 3D EM reconstruction. **A)** Left. Coomassie-stained SDS gel of the BBS2-7-9 complex purified by Strep-Tactin affinity purification. Flag-tagged human BBS9 (99.3 kDa) migrated below the 120 kDa molecular weight standard, and Strep-tagged human BBS7 (80.4 kDa) and HPC4-tagged human BBS2 (79.9 kDa) migrated near the 90 kDa molecular weight standard. The identity of these protein bands was confirmed by immunoblotting (right). **B)** Silver-stained SDS gel of the BBS2-7-9 complex purified as in panel A and separated by a glycerol gradient. The post-centrifugation fractions were loaded from the top (left) to the bottom (right). The fractions with a blue line (fraction numbers 10-13) were used for EM analysis. **C)** A representative image of a negatively-stained field of BBS2-7-9 complex particles (bar = 500 Å). **D)** Three orthogonal images of the 3D reconstruction of the BBS2-7-9 complex. (bar = 100 Å).

### 3D reconstruction of BBS2-7-9

We assessed the homogeneity of the complex by negative stain EM and found that despite the purity of the preparation, the particles were not sufficiently homogeneous for a high-resolution structural analysis by cryo-electron microscopy (Fig. 1C). As a result, we carried out a low-resolution 3D reconstruction using negative-stained EM images. The reconstruction (23 Å resolution) revealed the overall structure of the complex (Fig. 1D), which can be described as a flattened, triangular structure with a ∼ 200 Å height and ∼ 120 Å width at the base, with a small mass ∼ 40 Å in diameter protruding from one of the sides.

### XL-MS analysis of BBS2-7-9

In the absence of high-resolution cryo-EM data, we used the low-resolution EM envelope combined with cross-link mass spectrometry (XL-MS) to generate a molecular model of the BBS2-7-9 complex. XL-MS has become an effective tool to probe the structural architecture of protein complexes (32, 33). The crosslinks identified by XL-MS provide distance constraints that can be combined with EM reconstructions and other structural information to define the structures of protein complexes (34–39) (Fig. S1A). We treated the purified sample with increasing amounts of the crosslinker to determine the optimal concentration of crosslinker to use (Fig. S1B). Ultimately, the purified BBS2-7-9 was crosslinked with three different lysine-specific crosslinkers, disuccinimidyl suberate (DSS), disuccinimidyl glutarate (DSG) or Leiker reagent (40) and digested with proteases. The digests were analyzed by LC-MSMS, and crosslinked peptides were identified using the crosslink search engines xQuest (41) and pLink (42). Only high confidence hits that satisfied the screening criteria (see Experimental Procedures) were considered as true crosslinks. The coverage was nearly complete with 86 % (BBS2), 90 % (BBS7) and 87 % (BBS9) of all lysine residues reacting with the crosslinking reagents. Of those, 68 % (BBS2), 66 % (BBS7) and 78 % (BBS9) were involved in crosslinks while the rest formed monolinks (Table S4).

### Domain modeling

XL-MS generates three types of crosslinks that provide different structural information: crosslinks within individual domains of the BBSome subunits (intra-domain crosslinks), crosslinks between domains of the subunits (inter-domain crosslinks) and crosslinks between subunits (inter-protein crosslinks). The XL-MS analysis identified 51 intra-domain crosslinks (Fig. 2). These crosslinks were used to validate structural models of the individual domains of the core subunits. BBS2, 7 and 9 are homologous proteins that share a well-defined domain organization with N-terminal β-propeller, followed by coiled-coil, γ-adaptin ear (GAE), platform, and C-terminal α-helical domains (11). One atomic structure of the β-propeller domain of BBS9 has recently been reported (43), and the structures of homologous domains in other proteins have been solved (11, 19). Given these homologs, we reasoned that accurate structural models of each domain could be determined, and we generated homology models of each domain using the protein structure prediction server I-TASSER (44) (Fig. 2A, Table S5). The percentage of sequence modeled in these domains for BBS2, BBS7 and BBS9 was 95 %, 96 % and 96 %, respectively (Table S6).

**Figure 2.**
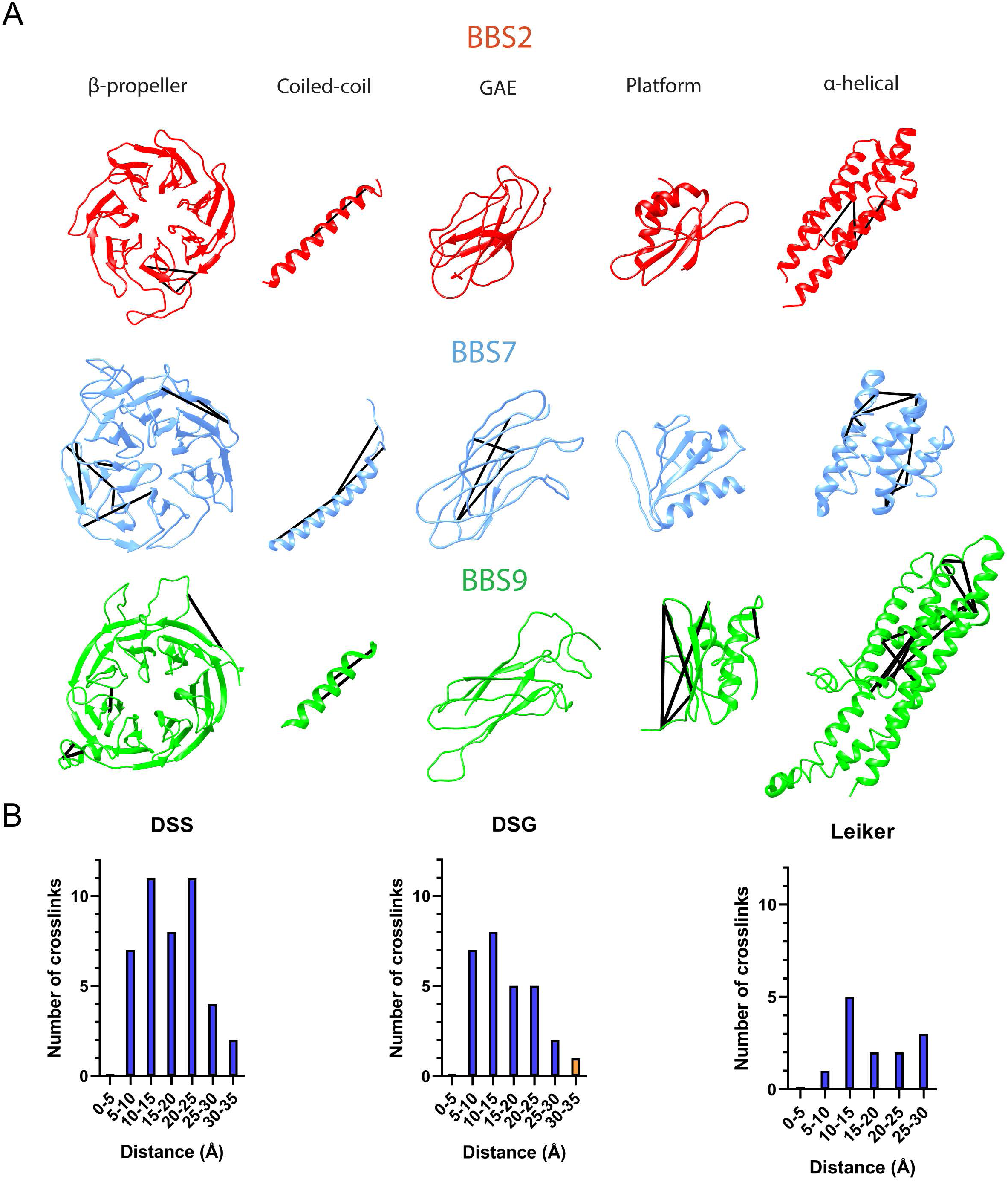
Domain structural models of the components of the BBS2-7-9 complex. **A)** Models of the domains of BBS2-7-9 complex components were created in I-TASSER except for the BBS9 β-propeller domain, which was from an X-ray crystal structure (PDB 4YD8). Intra-domain crosslinks were mapped onto the structures. **B)** Euclidean Cα distance distribution of the lysine crosslinks based on the domain structural models. All but one crosslink fit inside the expected distance constraints of 35 Å for DSS, 31 Å for DSG, and 33 Å for Leiker.

To assess the accuracy of the domain models, we mapped a list of all the theoretical lysine pair crosslinks onto the domain models and calculated Cα-Cα distances between the lysine residues. Considering movement of the protein backbone in flexible regions, the distance constraints were < 35 Å for DSS, < 31 Å for the DSG and < 33 Å for Leiker (36). We found that 21 % of all possible crosslinks in the domain models exceeded the 35 Å DSS distance constraint (Table S7). Yet when we mapped the experimentally determined intra-domain crosslink distances on the model structures, only one DSG crosslink of the 51 intra-domain crosslinks fell outside the distance constraints (Fig. 2B). This consistency between the structural models and the crosslinking distances validates the accuracy of the domain models for BBS2, 7 and 9 and the quality of the crosslinking data. These domain models were used as building blocks to assemble the BBS2-7-9 structure.

### BBS2 and BBS7 coiled-coil interaction

XL-MS identified 22 inter-protein crosslinks which were used to locate sites of interaction between subunits of BBS2-7-9 (Fig. 3A). Sixteen of the crosslinks were between BBS2 and BBS7 and six of these were in regions predicted to form a coiled-coil (residues 334-363 of BBS2 and 339-376 of BBS7). This high density of crosslinks supports the idea that BBS2 and BBS7 associate via a coiled-coil interaction in these regions. To determine if the coiled-coil interaction was parallel or anti-parallel, we modeled the coiled-coil region to both parallel and anti-parallel templates using the Foldit program (45) (Fig. S3B). When the cross-linking constraints were included in the modeling, the parallel template produced a compact coiled-coil structure with cross-link distances between 12-15 Å (Fig. 3B). In contrast, applying the cross-link constraints converted the anti-parallel template model into a parallel one, demonstrating that the cross-links were only compatible with a parallel model. The parallel model predicted that residues 334-363 of BBS2 participated in the coiled-coil with residues 339-363 of BBS7 and that residues 364-376 of BBS7 were not part of the coiled-coil (Fig. 3B). The interface between the two helices was populated by hydrophobic residues as expected for a coiled-coil interaction (Fig. 3C). Interestingly, no inter-protein crosslinks were found in the same coiled-coil region of BBS9, indicating that BBS9 does not form a coiled-coil interaction with BBS2 or BBS7. These data are consistent with pair-wise co-immunoprecipitation experiments that showed interactions between BBS2 and BBS7 as well as between BBS2 and BBS9 but not between BBS7 and BBS9 (23, 27).

**Figure 3.**
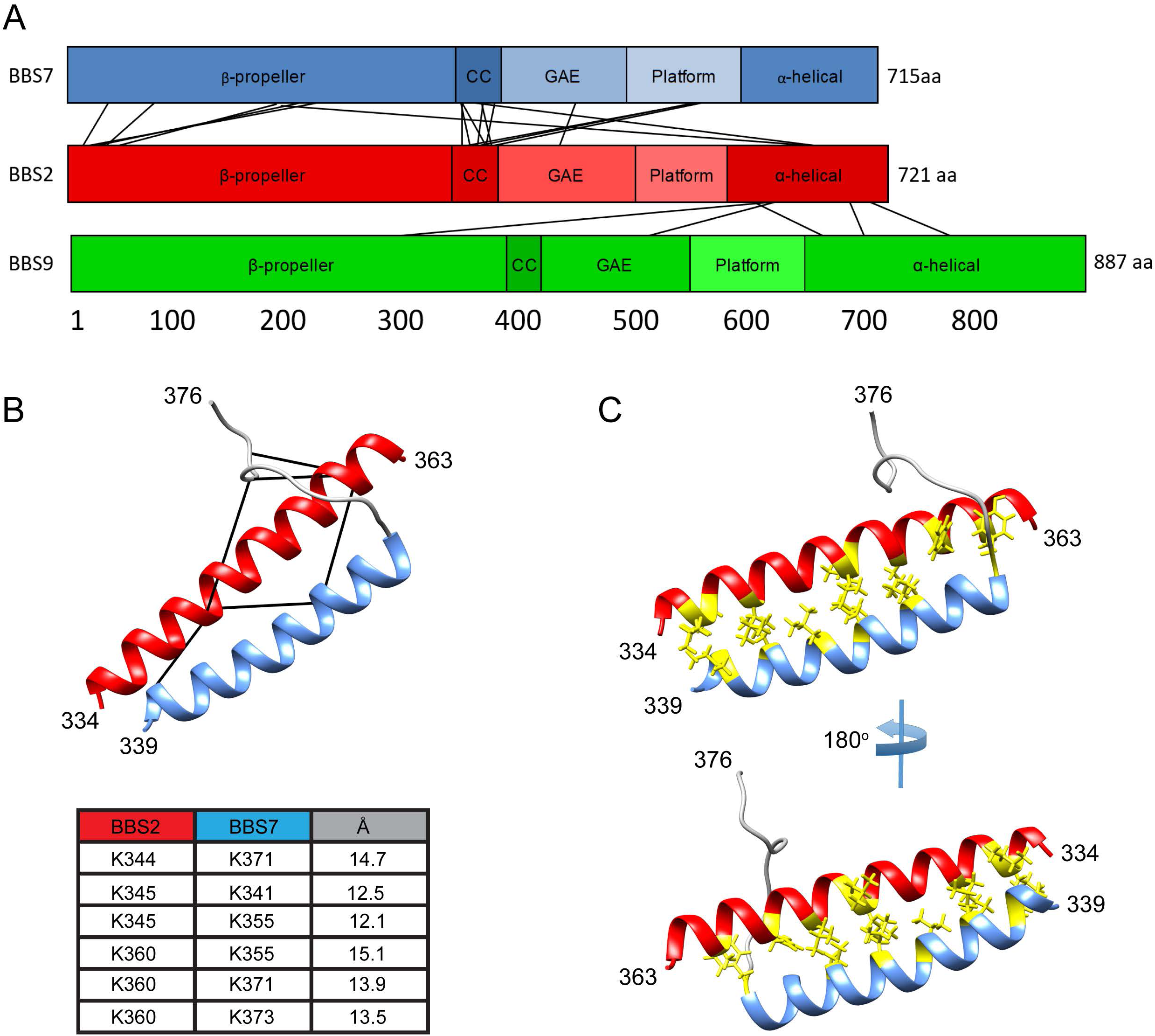
Coiled-coil interaction between BBS2 and BBS7. **A)** BBS2-7-9 complex inter-protein crosslinks determined by XL-MS. Six of these interlinks were found between the coiled-coil regions of BBS2 and BBS7. **B)** Structural model of the coiled-coil interaction between BBS2 and BBS7 with crosslinks mapped onto the model. The table lists the Euclidean Cα distances of the crosslinked lysines. The unstructured region of BBS7 (residues 364-376) is shown in gray. **C)** Two views of the BBS2-BBS7 coiled-coil interaction with the hydrophobic side chains at the interface shown in yellow.

### Structural Model of the BBSome core complex

To obtain a structural model of the entire BBS2-7-9 complex, we combined the BBS2/BBS7 coiled-coil model along with the domain models of the three BBS subunits and assembled them into the EM reconstruction using the 37 inter-domain crosslinks and the 22 inter-protein crosslinks identified by XL-MS. There were an additional five inter-domain crosslinks that mapped to unstructured regions between the modeled domains. These crosslinks could not be used for the integrated modeling. We employed the integrative modeling platform (IMP) (28), which takes structural data from the crosslinks and the EM density and converts them into spatial restraints that are combined into a scoring function to rank alternative models (46). IMP iteratively searches the configurational space to generate structural models that satisfy the spatial restraints, avoid steric clashes and retain sequence connectivity (46).

IMP iterations were run until the models converged to the point that they were producing similar structures. From these, the top 500 scoring models were compared in a distance matrix that calculates the root mean square deviation (RMSD) between the models (Fig. S2A). The matrix shows two clusters of models with related structures of 229 and 269 models each. Within each cluster, IMP calculates the localization density from all the models, which gives the overall shape of each protein in the complex. IMP also determines the global structural centroid for the cluster and selects the individual model whose centroid is nearest to this global centroid. The localization density and ribbon structure of the centroid model from cluster 1 is shown in Fig. 4A. Though the two clusters both fit the crosslinking and EM data comparably well, the first cluster satisfied more crosslinks than the second cluster, with 89.6 % and 85.4 % of crosslinks satisfied, respectively (Fig. 4B-C and Fig. S2B). The localization density of model 1 also fits the shape of the EM density well (Fig. 4D). The main differences in conformation between the two clusters is that in cluster 1, the β-propellers of BBS2 and BBS7 point out toward the short side of the EM density, whereas in cluster 2, they point out the long side. Otherwise, the inter-domain contacts between subunits in each cluster of models are the same (Fig. S2C).

**Figure 4.**
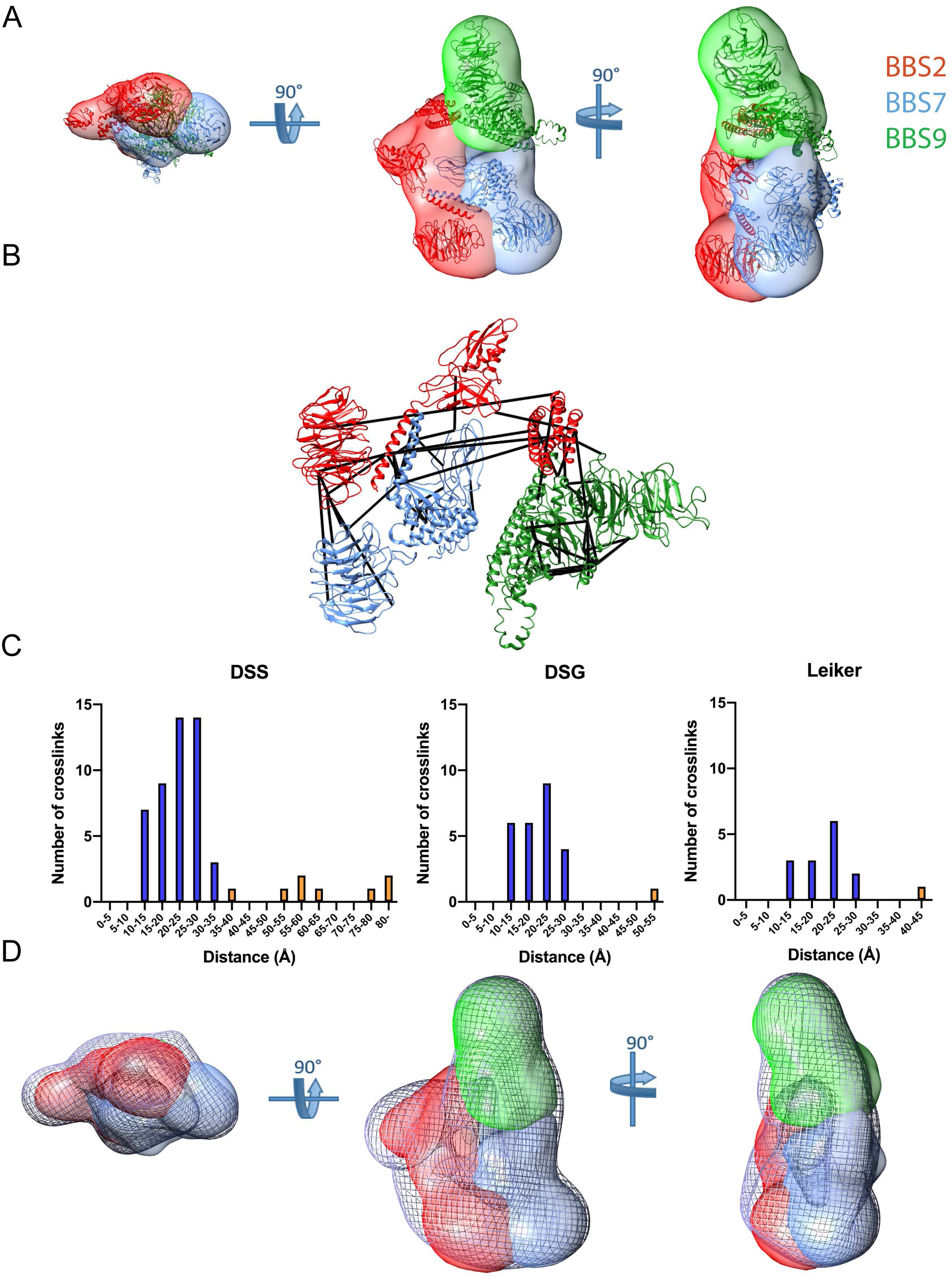
Structural model of the BBS2-7-9 complex. **A)** Three orthogonal views of the IMP localization densities computed for each subunit. The ribbon structure of the centroid model is depicted. **B)** Inter-domain and inter-protein crosslinks mapped onto the centroid model ribbon structure. **C)** Distributions of Euclidean Cα distances of the crosslinked lysines for each crosslinker. Blue bars represent crosslinks within the expected distance constraints and orange bars represent crosslinks outside the expected constraints (35 Å for DSS, 31 Å for DSG, and 33 Å for Leiker). **D)** The same views as in A docked in the EM 3D reconstruction of BBS2-7-9 complex. The reconstruction is depicted as a gray mesh surface.

An examination of the model structure shows BBS2 sitting between BBS7 and BBS9 and making close contact with both subunits (Fig. 4A). The coiled-coil is at the center of the interaction between BBS2 and BBS7, but their GAE domains are also in close proximity. The α-helical domain of BBS2 wraps around to contact the α-helical domain of BBS9, forming the primary point of contact between the two subunits. This proposed structure of the BBS2-7-9 complex is consistent with two previous studies showing binding of BBS2 to both BBS7 and BBS9 and no interaction of BBS7 with BBS9 (23, 27).

### Effects of BBS-linked mutations

Multiple point mutations in BBS2, BBS7 and BBS9 have been linked to Bardet-Biedl syndrome (30,43,47–54). To understand how these mutations might result in BBS, we measured the effect of 14 of these mutations (7 in BBS2, 6 in BBS7 and 1 in BBS9) on the formation of the BBS2-7-9 complex. We co-expressed BBS2, BBS7 and BBS9 in HEK-293T cells and measured the interaction of BBS7 and BBS9 with BBS2 by co-immunoprecipitation. In the case of BBS7, only the BBS2 L349W mutant showed a modest 30 % decrease in binding (Fig. 5A). This mutation is located in the BBS2/BBS7 coiled-coil (Fig. 3) and may weaken the coiled-coil interaction. In the case of BBS9, both the BBS2 R632P and the BBS9 G141R mutants showed a marked 70% reduction in binding. The decrease observed in the BBS9 G141R mutant was a result of lower cellular expression (Fig. 5B). Quantification of the BBS9 bands from cell lysates showed that expression of the G141R mutant was diminished by 76 % compared to the expression of wild-type BBS9. In contrast, expression of the BBS2 R632P mutant showed a slight 24 % decrease that was not statistically different from wild-type BBS2, indicating that the mutation did not significantly impair the expression of BBS2, but that it disrupted the interaction between BBS2 and BBS9. The position of the R632P mutation in the structural model suggests how the mutation would disrupt the interaction between BBS2 and BBS9. The R632P mutation is found within the α-helical domain of BBS2 and is in close proximity to α-helical domain of BBS9 (Fig. 5C). A proline substitution here would likely disrupt the formation of its helix and perturb the structure in this region, thereby interfering with the interaction between BBS2 and BBS9.

**Figure 5.**
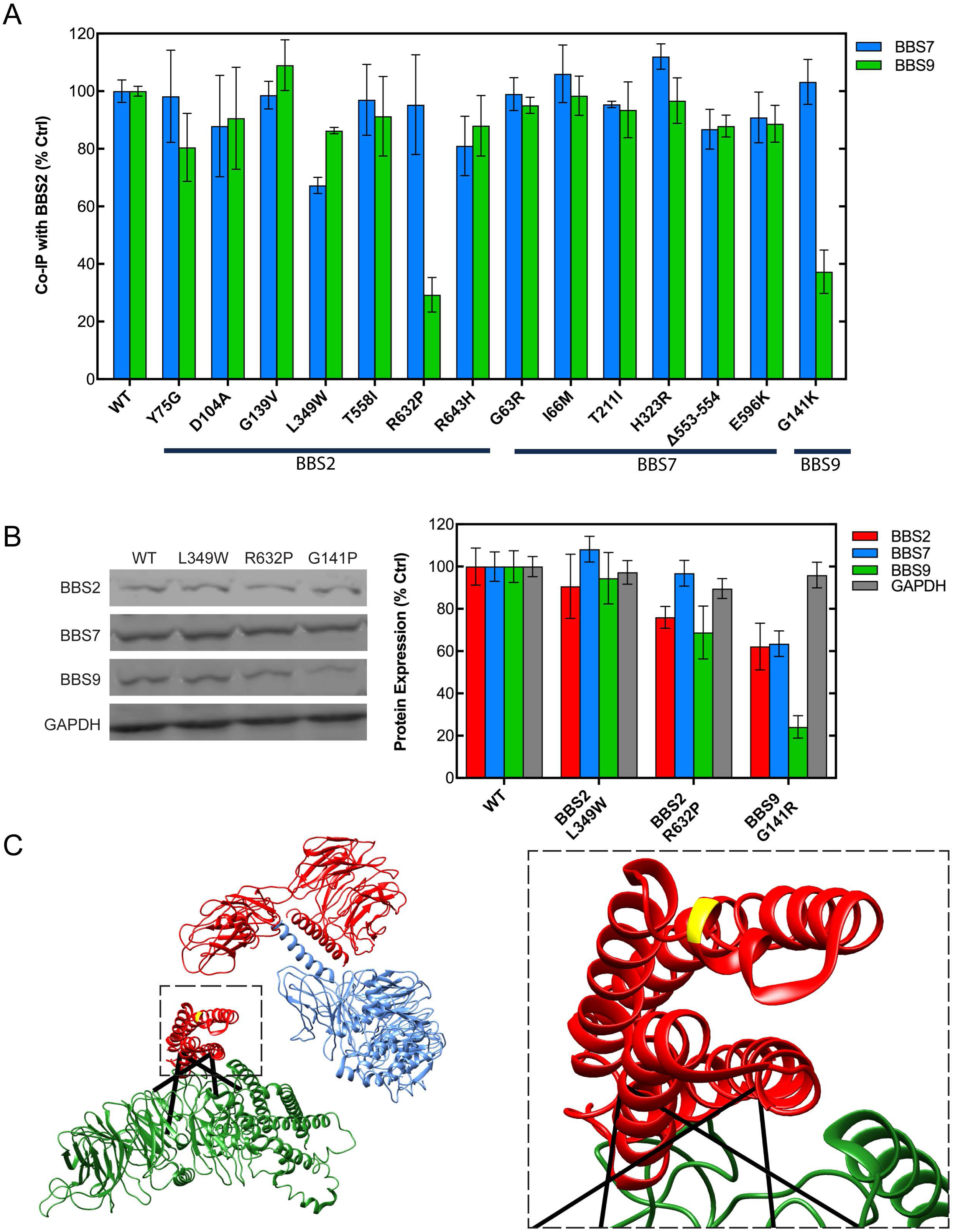
Effects of mutations on BBS2-7-9 complex assembly. **A)** The effects of BBS-linked mutations in BBS2, 7 and 9 on BBS2-7-9 complex formation were measured. HEK-293T cells were co-transfected with c-Myc-tagged wild type or mutant BBS2, flag-tagged wild type or mutant BBS7, and Flag-tagged wild type or mutant BBS9. BBS2 was immunoprecipitated, and BBS2, 7, and 9 were detected by immunoblotting. The graph shows the quantification of the BBS7 and BBS9 bands normalized to WT. Bars represent the average ± SEM of three experiments for each mutant and nine for WT. ** p < 0.01, *** p < 0.001. **B)** Expression levels of BBS2 L349W, R632P and BBS9 G141R mutants in cell lysates were determined by immunoblotting. Representative blots are shown and the graph gives the quantification of the BBS2, BBS7 and BBS9 bands normalized to WT. Bars represent the average ± SEM from six experiments. * p < 0.05, ** p < 0.01, *** p < 0.001. **C)** BBS2 R632P position. The BBS causing mutation BBS2 R632P (colored yellow) is located at the interface of BBS2 (red) and BBS9 (green). The zoomed in image shows the proximity of the BBS9 α-helical domain to BBS2 R632P. Lines indicate cross-links observed in this region.

## Discussion

To perform its essential role in ciliary transport, the eight subunits of the BBSome must be assembled into a functional complex. BBS2, BBS7 and BBS9 form a core complex that is an early intermediate in the assembly process (23). Our structural analysis of the BBS2-7-9 complex provides mechanistic insight into BBSome formation. BBS2 and BBS7 form a tight dimer principally via an extensive coiled-coil interaction involving residues 334-363 of BBS2 and residues 340-363 of BBS7. Previous studies indicate that BBS2 and BBS7 are brought together by chaperones, including the chaperonin-like BBS proteins (BBS6, BBS10 and BBS12) and the cytosolic chaperonin CCT (22, 23). BBS9 associates with the BBS2/BBS7 dimer through an interaction with the α-helical domain of BBS2. In contrast, BBS9 makes limited contacts with BBS7. Thus, the core complex is brought together by separate interactions of BBS2 with BBS7 and BBS9.

The roughly triangular structure of the core complex provides a suitable platform upon which the rest of the BBSome subunits can assemble (Fig. 6). BBS1 has been shown to interact strongly with BBS9 (27). Interestingly, BBS1 has a similar domain structure to BBS2, 7 and 9 with an N-terminal β-propeller followed by a coiled-coil and a GAE domain (11). Given the conserved coiled-coil of BBS1, it seems likely that BBS1 interacts with BBS9 via a coiled-coil interaction similar to that of BBS2 and BBS7. In the structural model, the coiled-coil region of BBS9 is exposed on a surface, far from BBS2 or BBS7, where it could form a coiled-coil with BBS1 without rearranging the complex (Fig. 6). BBS9 also interacts with BBS5 and BBS8, acting as a hub for BBSome assembly (23, 27). The core complex structure suggests multiple sites on BBS9 where BBS5 and BBS8 might interact. Both the β-propeller domain and the platform domain of BBS9 are completely accessible for interactions. In fact, the only region of BBS9 that is not accessible is the surface where BBS2 binds.

**Figure 6.**
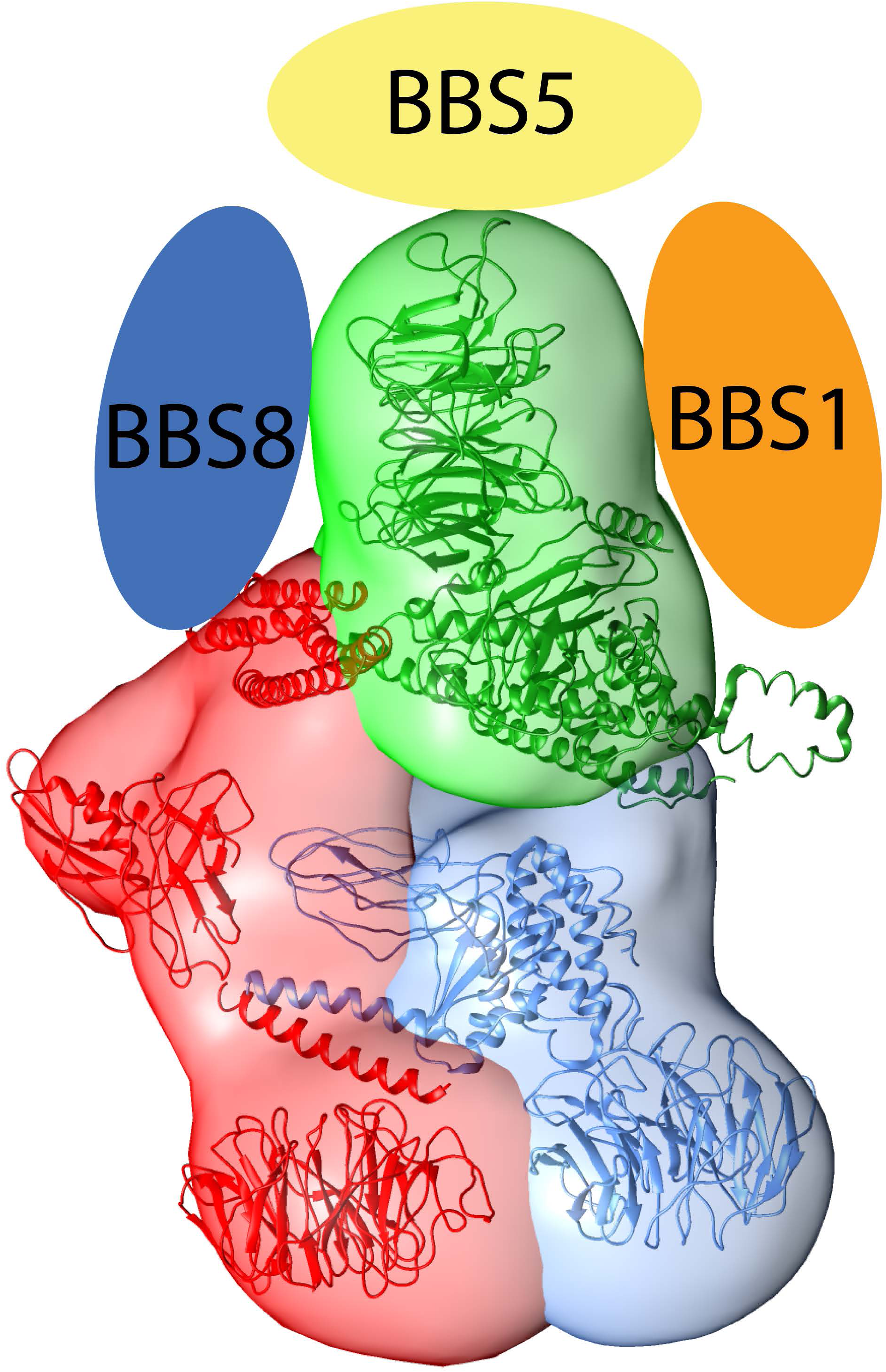
Cartoon model of the BBSome complex built around the structural model of BBS2-7-9 complex. BBS9 is the principal scaffolding protein of the BBSome that interacts with BBS1, BBS5 and BBS8 in addition to BBS2. The predicted coiled-coil region of BBS1 may bind BBS9 in an interaction homologous to the BBS2 and BBS7 coiled-coil. The nature of the interactions of BBS5 and BBS8 with BBS9 is not known.

The high accessibility of all three β-propeller domains in the core complex structure suggests that these domains might be sites of cargo binding in a manner analogous to the β-propellers of the α and β’ subunits of the COPI complex, which forms membrane vesicle coats for retrograde transport in the Golgi. These COPI β-propellers interact with membrane-associated cargos to bring them into vesicles (55–57). Similarly, the accessible β-propeller domains of the BBS2-7-9 complex may associate with membrane proteins for transport in the primary cilium.

The BBS2-7-9 complex structure provides insight into the molecular basis of BBS disease caused by the BBS2 R632P mutation. The interaction of this mutant with BBS9 is strongly inhibited, while its binding to BBS7 is not. The reason for the disruption can be seen in the predicted interaction surface between the α-helical domain of BBS2 with the α-helical domain of BBS9. Residues 628-635 of BBS2 are expected to form an α-helix that is located in the contact region between BBS2 and BBS9 (Fig. 5C). Three crosslinks between the α-helical domains of BBS2 and BBS9 (K609-K687, K673-K687, K691-K821) position these two domains together in the model and indicate that these domains closely interact (Fig. 5C). The R632P mutation would disrupt this helix of BBS2, which would in turn destabilize the interactions occurring at this interface. Thus, the R632P mutation likely causes BBS because of an inability of the BBSome core complex to assemble.

Previous work has shown that the G141R mutation disrupts the folding of the BBS9 β-propeller and inhibits the expression of the mutant protein (43). We also see decreased expression of the BBS9 G141R mutant that results in decreased binding of BBS9 to BBS2. Without BBS9, the BBSome cannot assemble and the disease ensues. The other mutants found in BBS2, 7 and 9 do not disrupt formation of the BBS2-7-9 complex, suggesting that they might contribute to BBS in some other way, perhaps by interfering with the association of the other BBSome proteins or BBSome cargos.

In summary, this structural analysis reveals the structural architecture of the BBS2-7-9 complex, consisting of BBS2, BBS7 and BBS9, and identifies key interactions that bring the components of the core complex together. Moreover, it provides a molecular basis for Bardet Biedl syndrome caused by the BBS2 R632P mutation. These studies show how structural information can inform our understanding of the BBSome and how its malfunction causes disease.

## Experimental Procedures

### Purification of the BBSome 2-7-9 Subcomplex

Human embryonic kidney (HEK)-293T cells cultured in 1:1 Dulbecco Modified Eagle media (DMEM) F12 with 10 % fetal bovine serum (FBS) in T-175 flasks were transfected at 80 % confluency with a transfection mixture consisting of purified plasmids of the pcS2+ vector (Addgene) containing human BBS2 with protein C peptide (HPC4) and c-Myc tags at the N-terminus, a pcDNA3.1 vector (Thermo Fisher Scientific) containing human BBS7 with a Flag-Strep tag at the N-terminus, a pcS2+ vector containing human BBS9 with a Flag tag at the N-terminus, and polyethylenimine (PEI). The DNA to PEI ratio was 1:3 by weight, and up to 90 μg of DNA was used per T-175 flask. The media was replenished with DMEM/FBS after 2-4 hours, and the cells were incubated for 48 additional hours before harvesting. Cells were lysed in 3 mL of extraction buffer (20 mM HEPES pH 7.5, 20 mM NaCl, 1 % NP-40, 0.5 mM PMSF, and 6 µL/mL protease inhibitors cocktail (Sigma P8340), 24 U/mL benzonase nuclease) per gram of cell pellet, and the lysate was cleared by centrifugation at 13,000 *g* for 15 minutes.

The BBS2-7-9 complex was purified at 4°C by affinity purification using 4 mL of Strep-Tactin resin (IBA Life Sciences) packed in a 2 cm diameter column and equilibrated with equilibration buffer (20 mM HEPES pH 7.5 and 20 mM NaCl). The HEK-293T cell lysate was loaded for 1 hour, washed with five column volumes of wash buffer (20 mM HEPES pH 7.5, 20 mM NaCl, and 0.05 % CHAPS), and then eluted with 2.5 column volumes of elution buffer (20 mM HEPES pH 7.5, 20 mM NaCl, 2.5 mM desthiobiotin, and 0.05 % CHAPS). The eluate was concentrated to ∼ 1 μg/μl using a 30 kDa cutoff filter (Millipore). The final protein concentration was determined by comparing the absorbance at 280 nm to a 2 μg/μl bovine serum albumin standard (Pierce) and correcting for minor buffer absorbance in a 2.5 μl nanodrop assay using a BioTek synergy H4 plate reader. The concentrated sample was flash frozen in liquid nitrogen and stored at −80°C. The purified core complex was analyzed by 10 % polyacrylamide SDS-gel electrophoresis and its purity was determined to be 80-90 % by staining with Coomassie Brilliant Blue (Thermo Fisher Scientific). The composition of the core complex was confirmed by immunoblotting with anti-Flag and anti-c-Myc antibodies. Coomassie gels and immunoblots were imaged using a LI-COR Odyssey infrared scanner.

Some samples were further affinity purified using an HPC4 antibody-conjugated resin (Roche), but yields from the HPC4 column were low, the improvements in purity were modest and the crosslinks identified were very similar to the Strep-Tactin purified complex. Therefore, we moved away from HPC4 purifications, but included crosslinking data from both purifications in the structural analysis.

### Crosslinking

The XL-MS analysis followed the protocol established by Leitner et al. (41). Approximately 200 μg of the BBS2-7-9 complex at 1 μg/μl was crosslinked in 25 mM HEPES pH 8.0, 100 mM KCl, and 325 μM of a 50 % mixture of H12/D12 disuccinimidyl suberate (DSS) or H6/D6 disuccinimidyl glutarate (DSG) (Creative Molecules) at 37°C for half an hour. The reaction was quenched by adding 50 mM ammonium bicarbonate to the crosslinked sample and incubating at 37°C for 15 minutes. The sample was dried by a vacuum concentrator, denatured in 100 mM Tris-HCl pH 8.5 and 8 M urea, reduced with 5 mM TCEP at 37°C for 30 minutes, and alkylated with 10 mM iodoacetamide at room temperature in the dark for 30 minutes. The sample was diluted with 150 mM ammonium bicarbonate to bring the urea concentration to 4 M, and proteins were digested with 4 μg of lysyl endopeptidase (Wako Laboratories, cleaves at the C-terminus of lysine) at a 1:50 enzyme to substrate ratio at 37°C for two hours. Subsequently, urea was diluted to 1 M, and proteins were further digested with trypsin (Wako Laboratories, cleaves at the C-terminus of lysine and arginine) at a 1:50 ratio at 37°C overnight. Peptide fragments were purified on a C18 column (Waters), dried, and reconstituted in 35 μL size exclusion chromatography (SEC) mobile phase (70:30:0.1 water:acetonitrile:TFA). Crosslinked peptide fragments were enriched by SEC using a Superdex Peptide PC 3.2/30 column on an AKTA pure system (GE Healthcare) at a flow rate of 50 μL/min. Fractions from the leading half of the elution peak were pooled, dried and resuspended in 2 % formic acid (41).

To increase the crosslink coverage, we also crosslinked with the Leiker trifunctional crosslinking reagent (40). We adapted the previously published protocol by first adding 50 μg of Leiker reagent to 200 μg of concentrated BBS2-7-9 complex in a final reaction volume of approximately 200 μL. The reaction proceeded for an hour and was quenched with 10 μL of 550 mM ammonium bicarbonate for 20 min. The crosslinked complex was then concentrated on a microcentrifuge filter (VWR 82031354) to < 20 μL and washed three times with 800 μL 8 M Urea, 20 mM methylamine, 100 mM Tris-Base, pH 8.5, reconcentrating to < 20 μL with each wash. The volume was brought up to 95 μL with the 8M urea solution, reduced with 5 mM TCEP, alkylated with 10 mM iodoacetamide, and digested by lysyl endopeptidase and trypsin as described above for the DSS and DSG crosslinking. The digested protein was then passed through the microcentrifuge filter and incubated with 40 μL of Pierce™ High Capacity Streptavidin Agarose (Thermo 20357) for two hours. The beads where then washed, the crosslinked peptides were eluted and the samples were prepared for mass spectrometry as described (40).

### Mass spectrometry

The enriched DSS and DSG crosslinked peptide samples were separated using a Thermo Fisher Scientific EASY-nLC 1000 liquid chromatograph system with a 15 cm Picofrit column (New Objective) packed with Reprosil-Pur C18-AQ of 3 μm particle size, 120 Å pore size and gradient of 5-95 % acetonitrile in 5 % DMSO and 0.1 % formic acid over 185 minutes and at a flow rate of 350 nL/min. The column was coupled via electrospray to an Orbitrap Velos Pro mass spectrometer. The resolution of MS1 was 30,000 over a scan range of 380-2000 m/z. Peptides with a charge state 3+ and greater were selected for HCD fragmentation at a normalized collision energy of 35 % with 3 steps of 10 % (stepped NEC) and a resolution of 7,500. Dynamic exclusion was enabled with a 10 ppm mass window and a one-minute time frame. Technical duplicates were run for each of three different DSS preparations and two DSG preparations.

The Leiker crosslinked samples were separated by LC-MS using an Easy-nLC 1200 system (Thermo Fisher Scientific) with a 75□μm×2 cm pre-column (3□μm C18 Acclaim PepMap 100 #164946) and a 75□μm×25 cm analytical column (2□μm C18 3□μm C18, Acclaim PepMap 100 #164946), using a gradient of buffer A (0.1% formic acid) and buffer B (80 % acetonitrile and 0.1 % formic acid) as follows: 0–6 % B for 2 min, 6–35 % B for 41 min, 35–100 % B for 10 min and 100 % B for 12 min at a flow rate of 300 nL/min. The column was coupled via electrospray to an Orbitrap Fusion Lumos mass spectrometer run in data-dependent mode with one scan at R = 60,000 (m/z = 350–2000), followed by ten HCD MS/MS microscans at R = 15,000 (first mass m/z = 110). The NCE was 27, with an isolation width of 2[m/z. The MS1 and MS2 scan AGC targets were 4e5 and 1e5 and the maximum injection time was 60 ms for both MS1 and MS2. Precursors of +1, +2, +7 or above, or unassigned charge states were rejected. Exclusion of isotopes was disabled. Dynamic exclusion was 30 s.

### XL-MS analysis

The xProphet (version 2.5.1)/xQuest (version 2.1.2) software pipeline (41) was used to identify the DSS and DSG crosslinked peptides, the specific lysine residues involved in the crosslink, and to evaluate the quality of each hit from the mass spectrometry data set. Tandem mass spectra for parent ions with a mass shift of 12.075321 Da for the DSS crosslinker and 6.04368 Da for the DSG crosslinker and a charge of +3 to +7 were classified as isotopic pairs and evaluated in ion-tag mode with the following parameters: 2 missed cleavages, 5-50 amino acid peptide length, carbamidomethyl fixed modification (57.02146 Da mass shift), oxidation variable modification (15.99491 Da mass shift), 138.0680796 (DSS) and 96.02059 (DSG) Da mass shift for intra-and inter-protein crosslinks, 156.0786442 (DSS) and 114.03115 (DSG) Da for –OH monolinks, 155.0964278 and 113.04713 for –NH_2_ monolinks, MS1 tolerance of 10 ppm, and MS2 tolerance of 0.2 Da for common ions and 0.3 Da for crosslink ions.

The peptide sequence database was created by xQuest based on the amino acid sequence of human BBS2, 7, and 9 (UniProt ID: Q9BXC9, Q8IW26, and Q3SYG4, respectively) and 9 common contaminant proteins (hHSPA1A, hHSPA1L, hHSPA8, hKRT1, hHSPA2, hTUBA1B, hTUBA1C, hHSPA6, hTUBA3C, UniProt ID: P0DMV8, P34931, P11142, P04264, P54652, P68363, Q9BQE3, P17066, P0DPH7, respectively). Spectra were searched against the database which covered all possible crosslink combinations of the BBSome core proteins, and any spectra that matched crosslinks in the database were counted and evaluated. Crosslink hits were screened with the following xQuest criteria: 10 % false discovery rate, > 10 % total ion counts, −4 to 7 ppm MS1 tolerance window, > 20 xQuest Id-Score, and > 4 fragmentation events per peptide. Any hit that met these parameters but scored below a decoy hit was not included in the crosslink list unless it was found in another run (Table S2). Those xQuest hits that satisfied these thresholds were inputted to the integrative modeling platform (IMP) for structural assessment (28).

All spectra were also analyzed with the pLink2 software suite (version 2.3.5). The same xQuest peptide sequence database was used and pLink added the sequences of other common contaminant proteins. The program was then run using the preset DSS and Leiker_clv linker settings and custom DSG (LinkerComposition: C(5)H(4)O(2), MonoComposition C(5)H(6)O(3)), DSG_heavy (C(5)H(−2)2H(6)O(2), C(5)2H(6)O(3)), and DSS_heavy (C(8)H(−2)2H(12)O(2), C(5)2H(12)O(3)) linker profiles using the same crosslink and monolink masses as described for xQuest. The spectra were then analyzed using conventional crosslinking (HCD) conditions, with trypsin set as the protease and up to 3 missed cleavages allowed. Peptides were selected with a mass between 600 and 6,000 Da and a length between 6 and 60 amino acids. The precursor and fragment tolerances were ± 20 ppm. The peptides were searched using carbamidomethyl (C) fixed modifications and phospho Y, T, S and oxidized M variable modifications. The results were filtered with a filter tolerance of ± 10 ppm and less than 5% FDR. (Table S3)

The residue numbers given in the xQuest and pLink output included amino-terminal tag lengths of 23 for BBS2, 16 for BBS7 and 11 for BBS9. These were subtracted to obtain the correct residue number of the crosslinked amino acids. Unique crosslinks were then sorted into intra-domain, intra-protein, and inter-protein data sets.

### Mutagenesis

The consequences of 14 BBS-linked mutations in BBS2, 7 and 9 on the formation of BBS2-7-9 were measured by co-immunoprecipitation. The mutations were introduced into N-terminally c-Myc-tagged BBS2 in pcS2+ vector, N-terminally Strep and Flag-tagged BBS7 in pcDNA3.1 vector and N-terminally Flag-tagged BBS9 also in pcS2+ vector using mutagenic PCR primers in a conventional PCR-based cloning protocol. All constructs were sequenced to confirm that the mutations were correct. Constructs were transfected into HEK-293T cells grown in 1:1 DMEM:F-12 media with 10 % FBS in 6-well plates at 80-90 % confluency using Lipofectamine 2000 according to the manufacturer’s protocol. Up to 3 μg of DNA was added to each well, and the relative DNA amounts were 1:1:1 for the BBS2, 7 and 9 constructs. Cells were fed with media three hours after transfection and incubated at 37°C for two days. Cells were washed with a PBS solution (12 mM phosphate pH 7.4, 137 mM NaCl, 3 mM KCl) and harvested in a PBS lysis buffer (PBS plus 6 uL/mL protease inhibitors cocktail (Sigma P8340), 0.6 mM PMSF, and 1 % NP-40) 48 hours after transfection. Lysed cells were triturated 8-10 times with a 25-gauge needle and syringe and cleared by centrifugation at 14,800 rpm in a Sorvall Legend Micro 21 microfuge for 10 minutes. The lysates were immunoprecipitated by incubating with an antibody to the c-Myc tag (Invitrogen 13-2500) on BBS2 as described above and immunoblotted with the c-Myc antibody for BBS2 and with the anti-Flag antibody (Sigma F3165) for BBS7 and BBS9 also as described above.

### Protein separation for electron microscopy (EM)

The BBS2-7-9 sample from the Strep affinity purification was concentrated to 1 μg/μl, and 150 μg of the sample was subjected to density gradient centrifugation to isolate the complexes. The sample was loaded onto a 10-30 % glycerol gradient in a separation buffer (20 mM HEPES pH 7.5, 100 mM KCl, 4 mM CaCl_2_, 0.3 mM PMSF, 0.3 % protease inhibitor cocktail, 3 mM DTT) and the ultracentrifugation was performed at 34,000 rpm in a SW55Ti rotor at 4 °C for 16 hours. Fractions were collected from the top to the bottom, analyzed by 10 % SDS-PAGE gels and silver stained. The fractions including pure BBS2-7-9 proteins were selected for the electron microscopy analysis.

### EM grid preparation and data collection

5 µl aliquots of the protein samples were applied to 300 mesh grids (Maxtaform Cu/Rh HR26) coated with a thin (∼ 8 nm) carbon layer and glow-discharged for 15 seconds. The grids were then stained (1 min) with 2 % uranyl acetate and air-dried before transmission EM analysis. Images were taken using Tecnai F20 transmission EM electron microscope operating at 200 kV with a 4k FEI Eagle CCD camera. Images were recorded at a sampling rate of 1.78 Å/px.

### Image processing and 3D reconstruction

Contrast transfer function was corrected using the CTFFIND3 program (58), which also calculated potential astigmatism. Micrographs with visible drift and astigmatism were discarded. 8806 single particles of the BBSome core complex were selected manually, extracted from micrographs, and normalized using the XMIPP software package (59). Three types of algorithms implemented in XMIPP were used to classify single images, CL2D (60) and Relion (61).

For 3D reconstruction, several initial models were tested in the first step of the 3D reconstruction procedure using EMAN software (62): artificial noise, blob, and a model created by a ‘common lines’ algorithm based on previously obtained 2D classes. Refinement was performed until the 3D reconstructions from these initial models converged to stable, similar 3D volumes. To obtain more structural detail, the 3D reconstruction from EMAN refinement was subjected to projection matching using XMIPP. Resolution of the final 3D models was estimated based on the FSC criterion (Fourier shell correlation) (63). The spatial frequency at 0.5 correlation was taken as the resolution of the model. Visualization of the 3D models was performed using USCF Chimera (64).

### Structural modeling

All three BBS2-7-9 proteins share the same domain organization with N-terminal β-propeller, coiled-coil, γ-adaptin ear (GAE), platform, and C-terminal alpha-helical domains. The secondary and tertiary structure of each domain was generated using the protein structure prediction server I-TASSER (44), and the accuracy of these domain models was confirmed using the intra-domain crosslinks from the XL-MS data. A crystal structure (PDB 4YD8) was used as a domain model of the β-propeller of BBS9 (43).

Sequence analysis indicated a coiled-coil interaction between BBS2 residues 334-363 with BBS7 residues 339-376. To assess this possibility, we employed homology modeling using the parallel coiled-coil from rat PAWR protein (chains A and B from PDB ID: 5fiy(65)) or the anti-parallel coiled-coil from the hantavirus nucleocapsid protein (2ic9) as templates to obtain a model structure for the BBS2/BBS7 coiled-coil. First, the coiled-coil heptad repeats were identified in the BBS2 and BBS7 sequences, revealing three clear repeats in each sequence (Fig. S3A). Similarly, three coiled-coil heptad repeats with ideal coiled-coil geometry were identified in the models and used as template structures. The program Foldit (45) was used to thread the BBS2 and BBS7 sequences into the templates. The triple heptad repeats of BBS2 and BBS7 were aligned with the heptad repeats of the templates and placed fully opposite and in register with one another. Foldit was then used to sample all side chain and backbone degrees of freedom in the resulting BBS2/BBS7 homology models. Restraints of 11.5 Å for DSS and 7.4 Å for DSG were applied between Nz atoms of the crosslinked lysine residues identified by XL-MS. With these restraints applied, the models were again allowed to sample all side chain and backbone degrees of freedom. The region of the homology model comprising BBS7 residues 364-376 did not fit the restraints well and so Foldit was used to unfold this region and refolded it using the aforementioned restraints. The result was an unstructured random coil for BBS7 residues 364-376 positioned outside the high confidence coiled-coil model for the rest of the structure.

The Integrative Modeling Platform (IMP) version 2.8.0 was used to predict how the individual BBSome core complex domains are oriented relative to each other. The domain models described previously were treated as rigid bodies. The parallel coiled-coil interaction of BBS2 and BBS7 was treated as a single rigid body. Linker regions between domains were approximated as 20 residue beads. Each of the three subunits was treated as super rigid bodies. Model restraints were generated from the crosslink data and from the EM reconstruction electron density. A weight of 120 was applied to the EM restraint. 3,200,000 models were generated in runs of 100,000 frames. The top 500 scoring models were then divided into two clusters and localization densities for each subunit were calculated for each cluster. Cluster precision was then gauged by calculating the root mean square fluctuation between models in the cluster and for each residue in each subunit (Fig. S2D). Model accuracy was determined by using the PDB file of the centroid model for all three subunits to calculate crosslink distances and to determine the fit within the EM density envelope.

### Experimental Design and Statistical Rationale

Three separate purifications of the BBS2-7-9 complex were crosslinked with DSS, two were crosslinked with DSG and four were crosslinked with Leiker reagent. All crosslinks that met the identification requirements were included in the IMP data. Binding experiments to measure the effects of BBS-linked mutations on BBS2-7-9 interactions were repeated three times in biologically independent assays to determine if differences from WT controls were significant. P-values were calculated using a 2-tailed T-test assuming a normal distribution.

## Supporting information

Crosslinks, spectra, modeling stats

## Footnote

This work was supported by Spanish Ministry of Economy and Innovation grant BFU2016-75984 (to JMV) and the US National Institutes of Health grant EY012287 (to BMW). The content is solely the responsibility of the authors and does not necessarily represent the official views of the National Institutes of Health. The authors declare that they have no conflicts of interest with the contents of this article.

## Data Availability

The raw MS spectra are deposited in the massIVE database (ftp://MSV000081472@massive.ucsd.edu) under the Project ID: MSV000081472 with username: WillardsonLab and password: willardsonbyu. The xQuest and pLink output files are available in Table S1.

## Supporting Information Figure Legends

**Figure S1.**
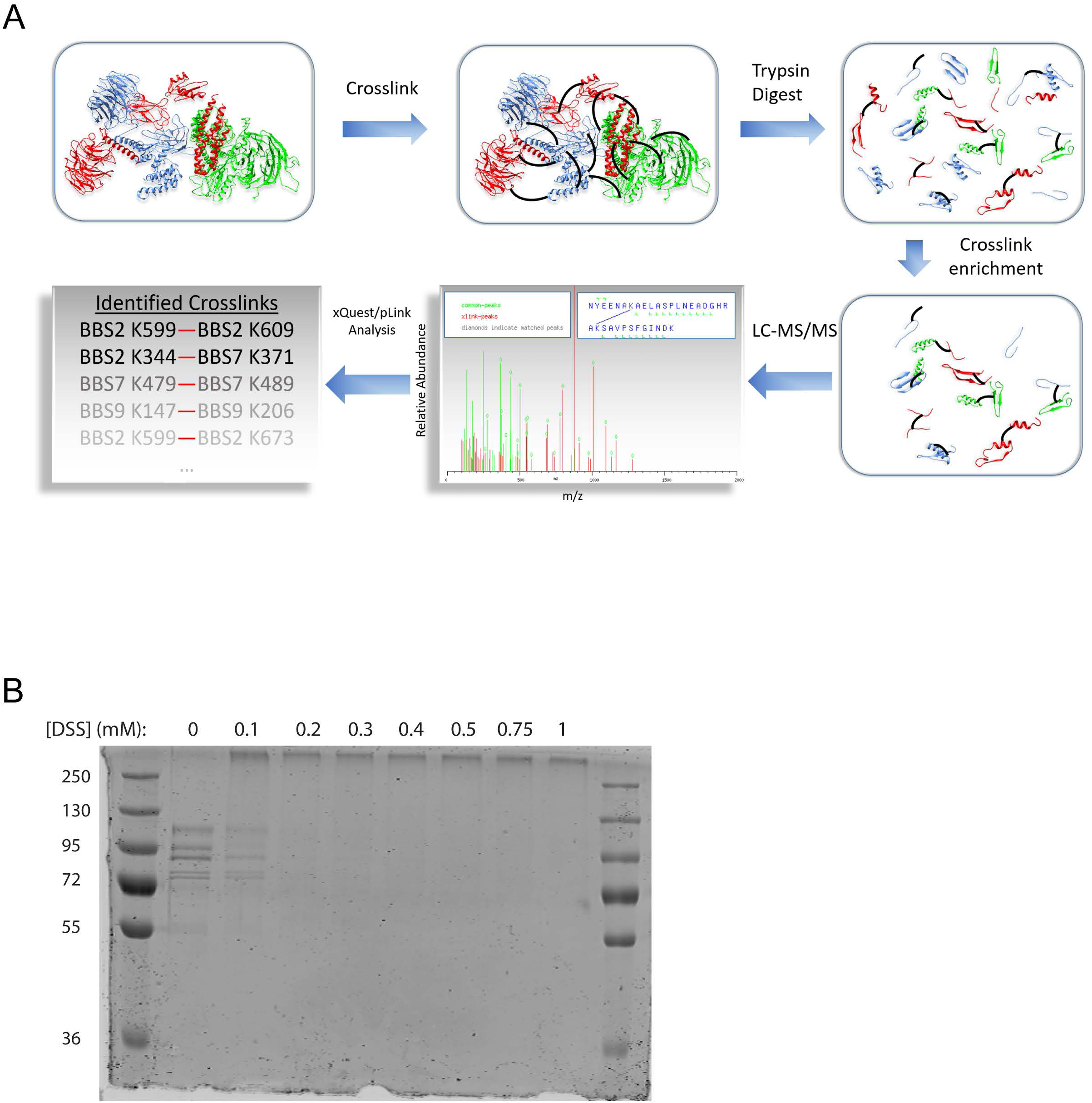
Workflow for the XL-MS experiments. **A)** The purified BBS2-7-9 complex was crosslinked and digested with proteases. Crosslinked peptides were enriched and analyzed by LCMS/MS. Cross-links were identified using the xQuest and pLink search engines. **B)** Coomassie-stained SDS gel of a titration of BBS2-7-9 complex treated with increasing amounts of DSS crosslinker. Bands at the top of the gel are multimeric crosslinked sample and lower bands are monomeric BBS2-7-9 subunits.

**Figure S2.**
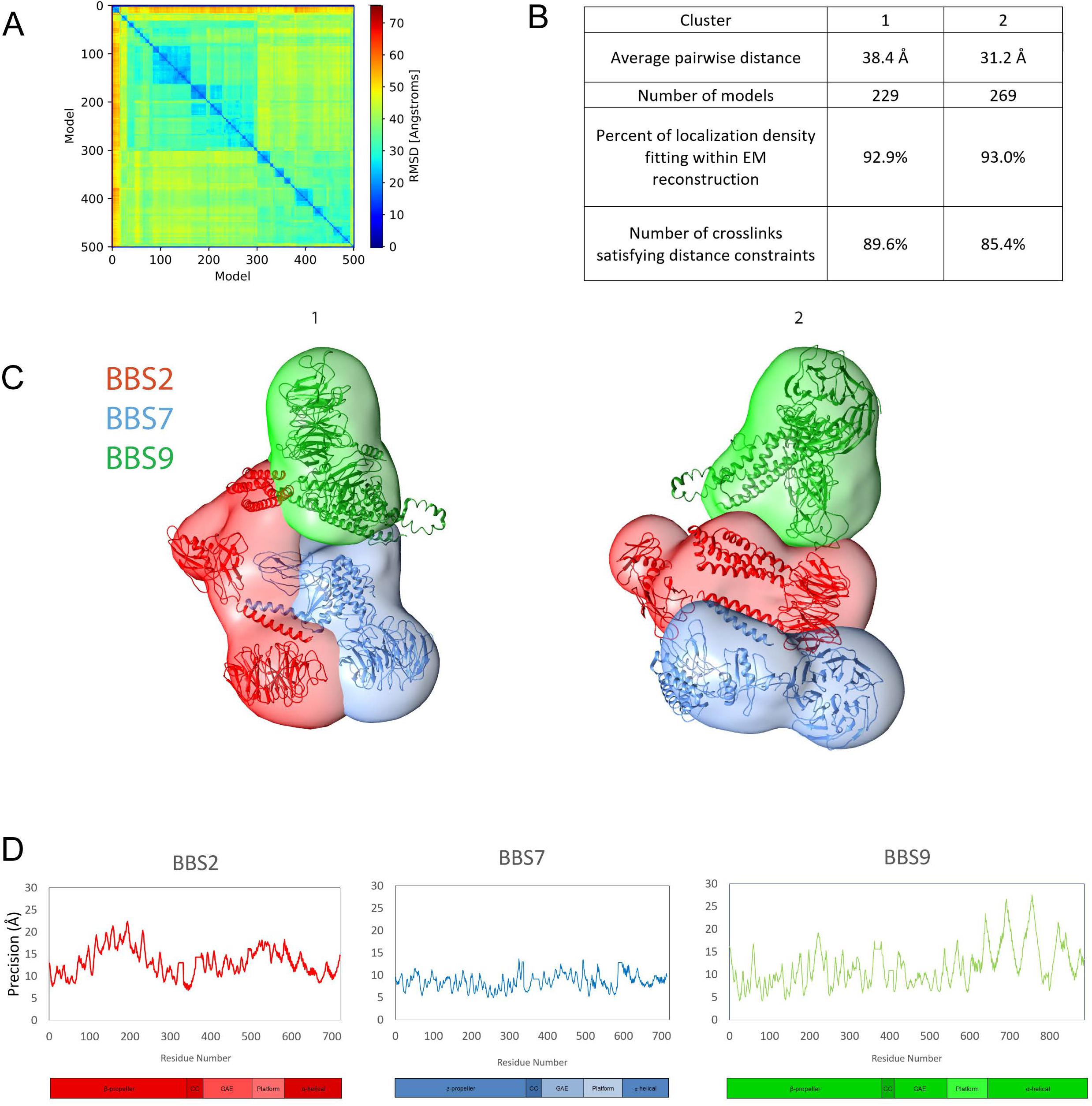
Precision of model clusters. **A)** Distance matrix of the RMSD between each of the top 500 scoring models. The three main clusters are outlined with a dashed line. **B)** Precision statistics for each cluster. **C)** Localization densities and centroid model ribbons structures for each cluster. **D)** Root mean squared fluctuation of each residue calculated for each subunit in cluster 3. Domain maps are shown below for reference.

**Figure S3.**
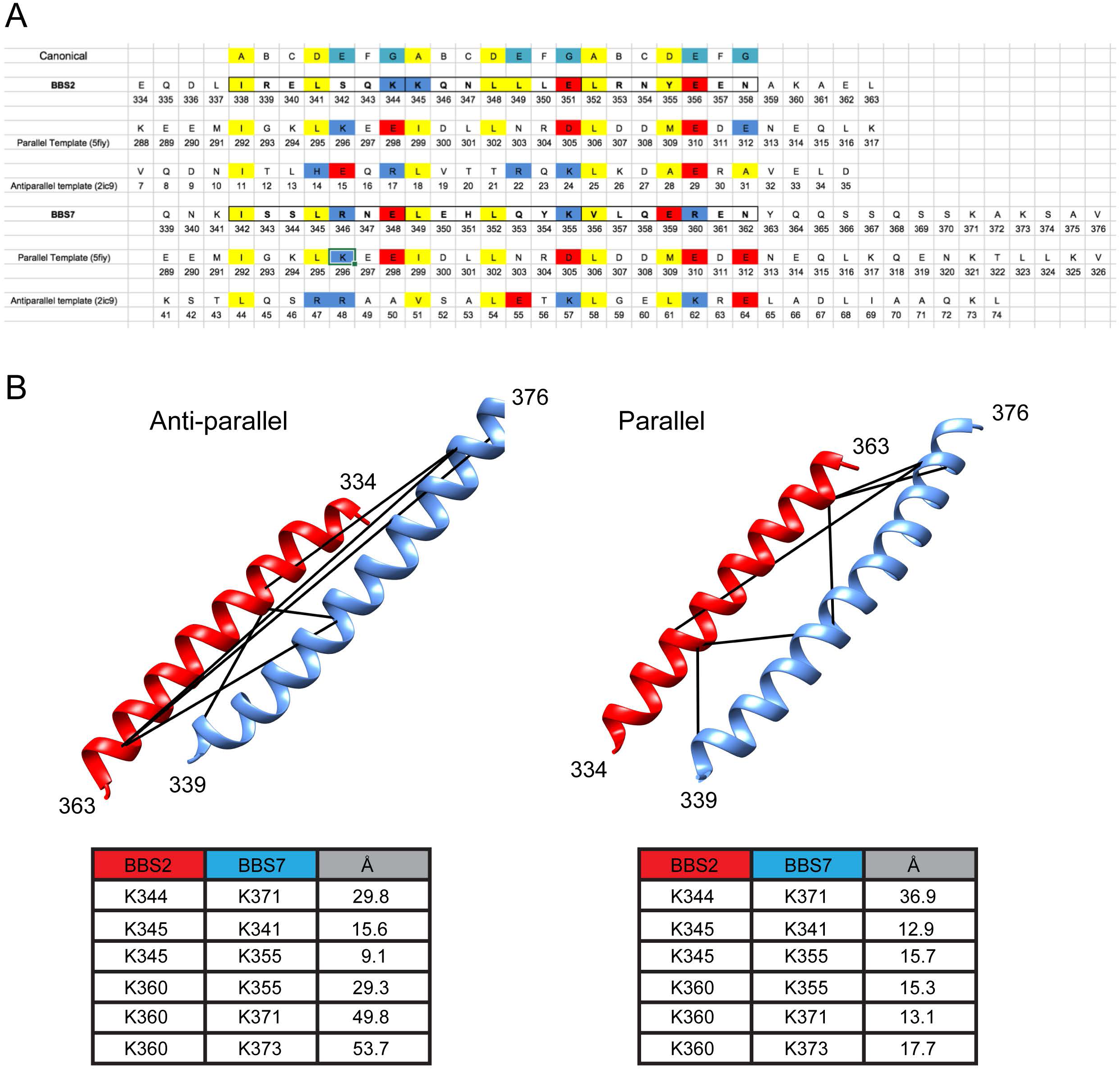
Orientation of the BBS2-BBS7 coiled coil. **A)** Sequence alignment of BBS2, BBS7 and canonical heptad repeat, together with the parallel and antiparallel template sequences. Residues are highlighted with yellow as hydrophobic, blue as acidic, and red as basic. **B)** Foldit models for the coiled-coil based on the anti-parallel and parallel templates. The Cα distances for each of the six crosslinks within the coiled-coil are listed below each model.

---

**Table S1. Master crosslink list.** The spreadsheet lists the unique crosslinks that met the xQuest and/or pLink criteria for use in the structural analysis. The protein and residue number of each crosslinked lysine is listed, followed by an X in each column representing the crosslinker used (DSS, DSG, or Leiker) and the search engine (xQuest or pLink).

**Table S2. Annotated xQuest spectra.** The spreadsheet gives the xQuest output from the analysis of the MSMS data of the five DSS or DSG crosslinking experiments. All crosslinks below a FDR of 10 % are shown.

**Table S3. Annotated pLink Spectra.** The spreadsheet gives the pLink output from the analysis of the MSMS data of the eight DSS, DSG, or Leiker crosslinking experiments. All crosslinks below a FDR of 5 % are shown.

**Table S4. Lysine crosslinking efficiency.** Each lysine residue is listed for BBS2, BBS7, and BBS9. Below the residue number is annotated whether there was no reaction with the linker observed (N), only a monolink reaction (M), only a crosslink reaction (X) or both a monolink and a crosslinking reaction. Total reaction types are listed for each subunit.

**Table S5. iTASSER model statistics.** Sequence for each domain of BBS2, BBS7, and BBS9 was submitted to the iTASSER server for homology modeling. The model quality statistics for each submission are listed. See https://zhanglab.ccmb.med.umich.edu/I-TASSER/annotation/ for a description of each statistic. The coiled-coil domains of BBS2 and BBS7 were modeled by Foldit and the β-propeller domain of BBS9 was solved by crystallography, so statistics on these models are not included.

**Table S6. Sequence modeling coverage.** The range of residues modeled in each domain is listed for each protein. In the final column, the percentage of all residues that were modeled as a rigid body in the Integrated Modeling Platform is indicated for each protein.

**Table S7. Theoretical Intra-domain Crosslinks.** The number of lysine residues in each domain is listed in the first column, followed by the number of possible _n_C_2_ combinations of lysine pairs. The Euclidean Cα distances for each lysine pair was calculated in chimera. The number of these theoretical crosslinks exceeding 35 Å is listed in the third column, followed by the percentage of all lysine pairs that exceed the 35 Å distance constraint.

